# Causal evidence supporting the proposal that dopamine transients function as a *temporal difference* prediction error

**DOI:** 10.1101/520965

**Authors:** Etienne JP Maes, Melissa J Sharpe, Matthew P.H. Gardner, Chun Yun Chang, Geoffrey Schoenbaum, Mihaela D. Iordanova

**Affiliations:** Department of Psychology/Centre for Studies in Behavioural Neurobiology, Concordia University, Montreal, H4B 1R6; Department of Psychology, University of California, Los Angeles, CA 90065.; NIDA Intramural Research Program, Baltimore, MD 21224; Departments of Anatomy & Neurobiology and Psychiatry, University of Maryland School of Medicine, Baltimore, MD 21201; Solomon H. Snyder Department of Neuroscience, The Johns Hopkins University, Baltimore, MD 21287

**Author notes:** senior corresponding authors, (GS), (MDI).

## Abstract

Reward-evoked dopamine is well-established as a prediction error. However the central tenet of temporal difference accounts – that similar transients evoked by reward-predictive cues also function as errors – remains untested. To address this, we used two phenomena, second-order conditioning and blocking, in order to examine the role of dopamine in prediction error versus reward prediction. We show that optogenetically-shunting dopamine activity at the start of a reward-predicting cue prevents second-order conditioning without affecting blocking. These results support temporal difference accounts by providing causal evidence that *cue-evoked* dopamine transients function as prediction errors.

---

One of the most fundamental questions in neuroscience concerns how associative learning is implemented in the brain. Key to most implementations is the concept of a prediction error – a teaching signal that supports learning when reality fails to match predictions ^1^. The greater the error, the greater the learning. In computational accounts, these errors are calculated by the method of temporal difference ^2,3^, in which time (t) is divided into states, each containing a value prediction (V) derived from past experience that is the basis of a rolling prediction error. This temporal difference prediction error (δ) is the difference between successive value states. The most famous of these is temporal difference reinforcement learning TDRL, ^3^, whose prediction error:

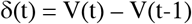

has been mapped onto transient, millisecond resolution changes in dopamine neuron firing that occur during reward learning ^4^.

While this mapping has been one of the signature success stories of modern neuroscience, one pillar of this account that has not been well-tested is that the transient increase in firing evoked by a reward-predicting cue is a *temporal difference* error, propogated back from the reward and functioning to support learning about predictors of that cue. Evidence for true, gradual backpropogation of this signal is sparse, as is evidence that it exhibits signature features that define the error at the time of reward, such as suppression on omission of the cue when it has been predicted by an earlier cue, and transfer back to such earlier predictors (in the absence of the primary reward itself). Further, there is little or no causal evidence that cue-evoked dopamine serves as an error signal to support learning. Indeed, the cue-evoked signal is often described as if it encodes the cue’s significance or value derived from its prediction of future reward. Such language is imprecise, leading at best to confusion about the theorized unitary function of the dopamine transient and at worst to a true dichotomization of the function of cue-versus reward-evoked activity. This situation is especially curious, since the appearance of the dopamine transient in response to reward-predictive cues is a lynchpin of the argument that the dopamine neurons signal a *temporal difference* error ^1^.

So does the cue-evoked dopamine transient reflect a prediction error or is it a reward prediction? A logical way to address this question is to test, using second-order conditioning ^5^, whether optogenetic blockade or shunting of dopamine activity at the start of a reward-predictive cue prevents learning about this cue in the same way that optogenetic shunting of dopamine activity at reward delivery prevents learning about reward ^6^. If this signal is a temporal difference error, δ(t_cue_) in the terms of the above equation, then blocking it will prevent such learning; this is illustrated in the top row of Figure 1 (and Figure S1), which shows an experimental design for second-order conditioning and computational modeling of the effect of eliminating δ(t_cue_). However, while this seems at first like a conclusive experiment, it is not, since the same effect is obtained by eliminating the cue’s significance or ability to predict reward for the purposes of calculating the prediction error (Figure 1, also Figure S1). This occurs because the cue’s ability to predict reward is the source of V(t_cue_), which is the basis of the cue-evoked prediction error. Therefore, if when you shunt the transient you are eliminating the cue-evoked prediction, you would also eliminate the cue-evoked prediction error as a consequence. As a result, the disruption of second-order conditioning by shunting of the dopamine transient would show that this signal is necessary for learning, but would fail to distinguish whether it is a prediction error or a reward prediction.

**Figure 1:**
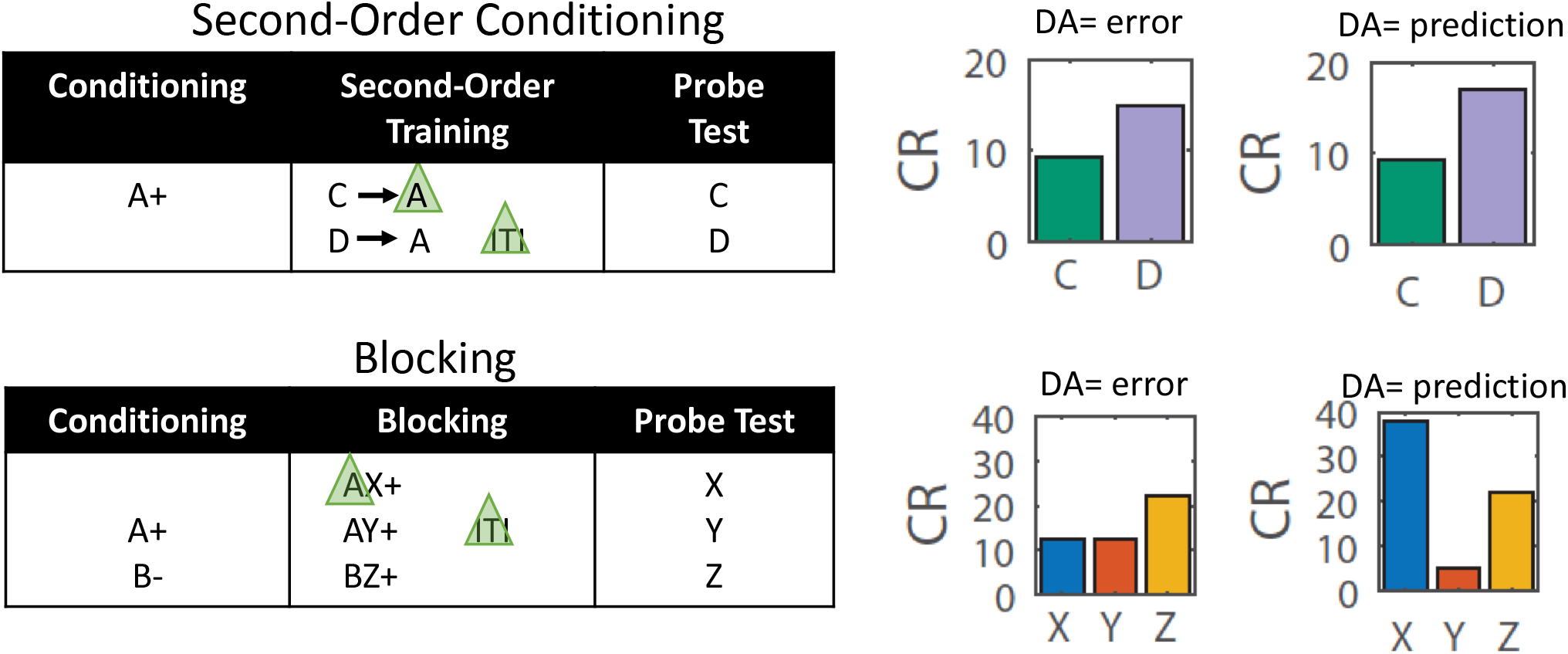
Modeling Results. Experimental designs for second-order conditioning (top row) and blocking (bottom row), along with bar graphs modeling the predicted results of shunting of the dopamine transient at the start of the reward-predictive cue, A, in each procedure. Green triangles indicate light delivery to shunt dopamine transients at the start of the cue in TH-Cre rats expressing halorhodopsin in VTA neurons. The left column of bar graphs shows modeled results under the hypothesis that the cue-evoked dopamine transient signals a prediction error; the right column shows them under the hypothesis that it signals a reward prediction. Elimination of either signal would impair second-order conditioning (top graphs, C versus D), but only elimination of a prediction would affect blocking (bottom graphs, X vs Y/Z). Note the output of the classic TDRL model was converted from V to conditioned responding (CR) to better reflect the behavioral output actually measured in our experiments (see methods for details). See figure S1 for modeling of behavior in the full experiments, culminating in these displays.

This confound can be resolved by combining the above experiment with an assessment of the effects of the same manipulation on blocking ^7^. Blocking refers to the ability of a cue to prevent or block other cues from becoming associated with the reward it predicts; blocking is thought to reflect the reduction in prediction error at the time of reward, δ(t_rew_), caused by the cue’s reward prediction, which becomes the prediction immediately before reward delivery (V(t_rew_ −1)). If the cue-evoked dopamine transient is carrying that prediction, then optogenetically-shunting it should diminish or prevent blocking, because in that case the reward would still evoke a prediction error; this is illustrated in the bottom row of Figure 1 (and Figure S1), which shows an experimental design for blocking and computational modeling of the effect of eliminating V(t rew −1). On the other hand, if the cue-evoked dopamine signal reflects only the actual prediction error occurring at the start of the cue, δ(t_cue_), then its removal should have no impact on blocking (Figure 1, also Figure S1).

Armed with these contrasting computationally-validated predictions, we set out to test them using a within-subjects version of the designs shown in Figure 1 (shown in Figure S1). Sixteen Long-Evans transgenic rats expressing Cre recombinase under control of the tyrosine hydroxylase promoter (Th-cre+/-) served as subjects. Four weeks prior to the start of testing, the rats underwent surgery to infuse a cre-dependent viral vector carrying halorhodopsin (AAV5-EF1α-DIO, eNpHR3.0-eYFP) into the ventral tegmengtal area (VTA) bilaterally and to implant optical fibers targeting this region. Histological analyses confirmed viral expression and fiber tip localization (Figure S2). Rats were food restricted immediately prior to the start of testing and then trained to associate a visual cue, A, with reward. The blocking experiment was run in the same rats before the SOC experiment to ensure that any effect of non-reinforcement during SOC training does not influence blocking. To align with the logic of our modeling, we present the SOC data first here.

Second-order training consisted of 2 sessions in which the previously-conditioned 13s visual cue, A, was presented without reward, preceded on each trial by one of two 10s novel auditory cues, C or D. On C→A trials, continuous laser light (532nm, 18-20mW output, Shanghai Laser & Optics Century Co., Ltd) was delivered into the VTA for 2.5s, starting 0.5s prior to the onset of A in order to disrupt any dopamine transient normally occurring at the start of reward-predicting cue, in this case, A. We model this as shunting, since we have found that similar patterns disrupt learning from positive errors without inducing aversion or learning from negative errors ^6,8,9^. On D→A trials, the same light pattern was delivered during the intertrial interval, 120-180s after termination of A, to serve as a control. Following this training, rats underwent probe testing, in which C and D were presented alone and without reward. The results are illustrated in Figure 2, with supporting statistics described in the caption. There was no difference in responding on C→A versus D→A trials, indicating that the optogenetic manipulation did not deter responding during second-order conditioning. However in the probe test, responding to C was significantly lower than responding to D, indicating that light delivery at the start of A prevented second-order conditioning of C.

**Figure 2:**
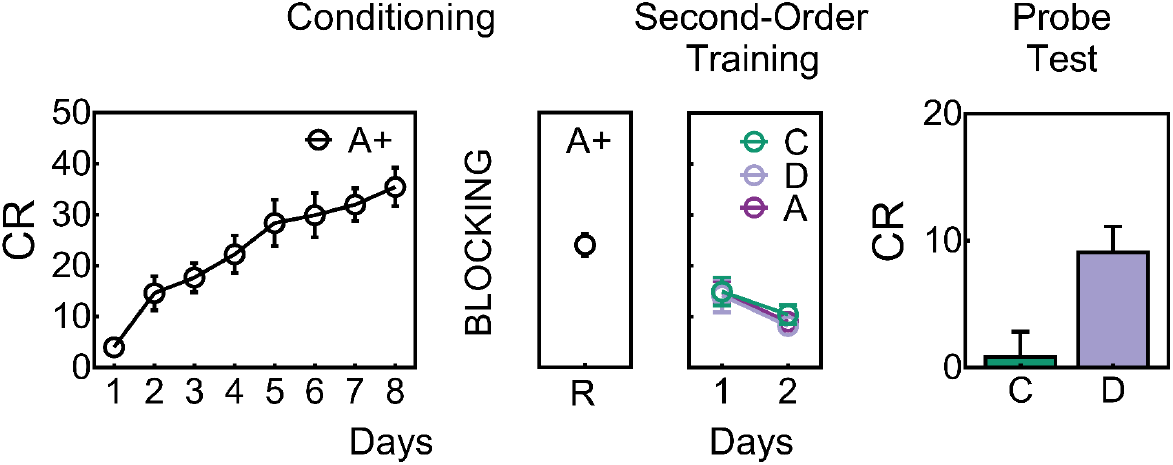
The cue-evoked dopamine transient is necessary for second-order conditioning. Behavioural responding during A increased during Conditioning and following Blocking during reminder (R) training (see online methods and Fig S2 for statistics). Responding to C (i.e., C→A trials) and D (i.e., D→A trials) did not differ (see online methods and Fig S2 for statistics) during second-order training when shunting of VTA transients took place at the start of the reward-predictive cue, A (as illustrated in Figure 1). On Test, responding to C was lower compared to D (t_15_=2.2, p=0.04), showing that inhibition of the VTA DA signal at the start of A prevented A from supporting second-order conditioning to C whereas identical inhibition during the ITI left learning to D intact. R, reminder training post-blocking. CR or conditioned responding is percent time spent in the magazine during the last 5s of the cue.

Blocking consisted of 4 sessions in which the previously-conditioned, 13s visual cue, A, was presented in compound with two novel 10s auditory cues, X and Y, followed by reward. On AX trials, continuous laser light (532nm, 18-20mW output, Shanghai Laser & Optics Century Co., Ltd) was delivered into the VTA for 2.5s, starting 0.5s prior to the onset of A; on AY trials, the same light pattern was delivered during the intertrial interval, 120-180s after termination of the compound. In addition, as a positive control for learning about a compound cue, the rats also received presentations of two additional cues, B and Z, followed by the same reward. B was a visual cue, which received four nonreinforced exposures daily across the last four days of conditoning (see Figure S1), to prevent any potential disruptive effects of unconditioned orienting to B during conditioning of the BZ compound. Z was a third novel auditory cue. Following this training, rats underwent probe testing, in which X, Y, and Z were presented alone and without reward. The results are illustrated in Figure 3, with supporting statistics described in the caption. During blocking, responding to BZ was lower at the start of training but reached that of the AX and AY compounds by the end of training. There were no differences in responding on AX versus AY trials across blocking, indicating that the optogenetic manipulation did not deter responding. In the probe test, responding to the positive control cue, Z, was significantly higher than responding to two cues blocked cues, X and Y, indicating that pretraining of A blocked learning for these two cues. Further, there was no difference in responding to X and Y, indicating that light delivery at the start of A on AX trials had no effect on blocking.

**Figure 3:**
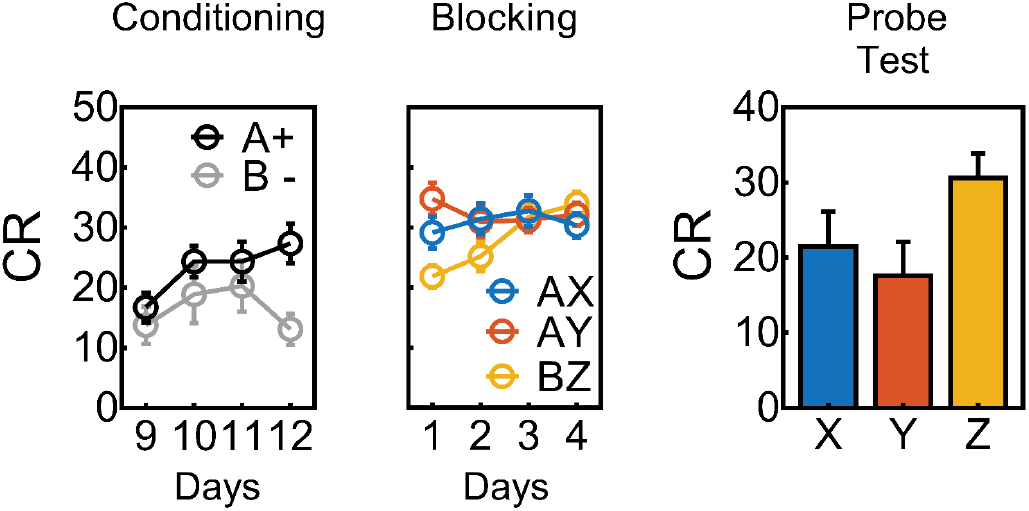
The cue-evoked dopamine transient is not necessary for blocking. Behavioural responding during conditioning was greater to the reinforced A compared to the non-reinforced B (see online methods and Fig S3 for statistics). During Blocking responding to the control compound (BZ) increased to the level of the blocking compounds (AX and AY); shunting of the VTA DA transient took place at the start of the reward-predicting cue, A (as illustrated in Figure 1), yet responding to AX and AY was similar on each day (see online methods and Fig S3 for statistics). On Test, the rats showed a blocking effect: there was higher level of responding to the control cue (Z) compared to the blocked cues (X and Y) (Figure 2c, F_1,15_=6.2, CI {1.03:0.08}) on the first trial pooled across both tests. There was no effect of VTA DA inhibition on blocking as responding to the blocked cues (X vs. Y) did not differ (F<1, CI{-0.60:1.06}). CR or conditioned responding is percent time spent in the magazine during the last 5s of the cue.

Our results show that shunting the VTA DA signal at the start of a reward predicting cue disrupted second-order conditioning but left blocking intact. This differential effect provides evidence that light artefacts from optical stimulation do not serving to hinder processing of A. If it did, then, we would see a disupriton of the blocking effect as well (i.e., learning about X). These data are important because they they provide clear and concise evidence that transient increases in the firing of dopamine neurons at the start of reward-predictive cues function as prediction errors to support associative learning in much the same way that reward-evoked changes have been shown to do. As noted in our introduction, this demonstration is important because the proposal that the cue-evoked dopamine transient is a prediction error is the lynchpin of the hypothesis that dopaminergic error signals integrate information about future events, thereby providing a temporal difference error. Further, by showing that cue-evoked firing of dopamine neurons is necessary for second-order conditioning, our results provide strong support for this idea, while at the same time ruling out alternative proposals that this signal reflects the actual associative significance of the cue with respect to predicting reward.

Importantly, our findings are agnostic with regard to the nature of the information in the temporal difference signal or the specific type of learning that it supports. This is a noteworthy caveat, since temporal difference errors can be limited to representing information about value ^2,3^ or they can be construed more broadly as representing errors in predicting other value-neutral information ^2,10^. Recent studies using sensory preconditioning and reinforcer devaluation provide evidence that dopamine transients support learning that is orthogonal to value and in line with the latter account ^9-11^. Here we used second-order conditioning, which has been proposed to rely on an associative structure that bypasses the representation of the outcome and links a stimulus and a response ^12^. Sensory pre-conditoning, by contrast, is supported by the association of two neutral stimuli, leaving no opportunity for direct links with a reward-based response. That dopamine transients in VTA are now implicated in supporting both forms of learning clearly supports a much broader role for these signals in driving associative learning than is envisioned by current dogma, and further suggests that perhaps the content of the learning supported is heavily determined by the learning conditions. Finally, it is worth noting that while our data support a role for VTA DA in prediction error within the context of Pavlovian associative learning in reward, they do not preclude other dopamine neurons broadcasting other content at target sites throughout the brain ^14^.

## Methods

Methods and any associated references are available in the online version of the paper.

## Acknowledgements

This work was supported by the Intramural Research Program at the National Institute on Drug Abuse; the Canada Research Chairs program (to MDI); a NSERC Discovery Grant (to MDI); a NSERC Undergraduate Student Research Award (to EJPM). The opinions expressed in this article are the authors’ own and do not reflect the view of the NIH/DHHS. The authors have no conflicts of interest to report.

## Author Contributions

EJPM, MJS, GS, and MDI conceived and designed the experiments, EJPM and MJS collected the behavioral data, CYC supervised the immunohistological verification of expression and fiber placement, and MPHG conducted the computational modeling. MPS and MDI analyzed the data, and GS and MDI interpreted the data and wrote the manuscript with input from the other authors.

## Supplemental Figures

**Figure S1:**
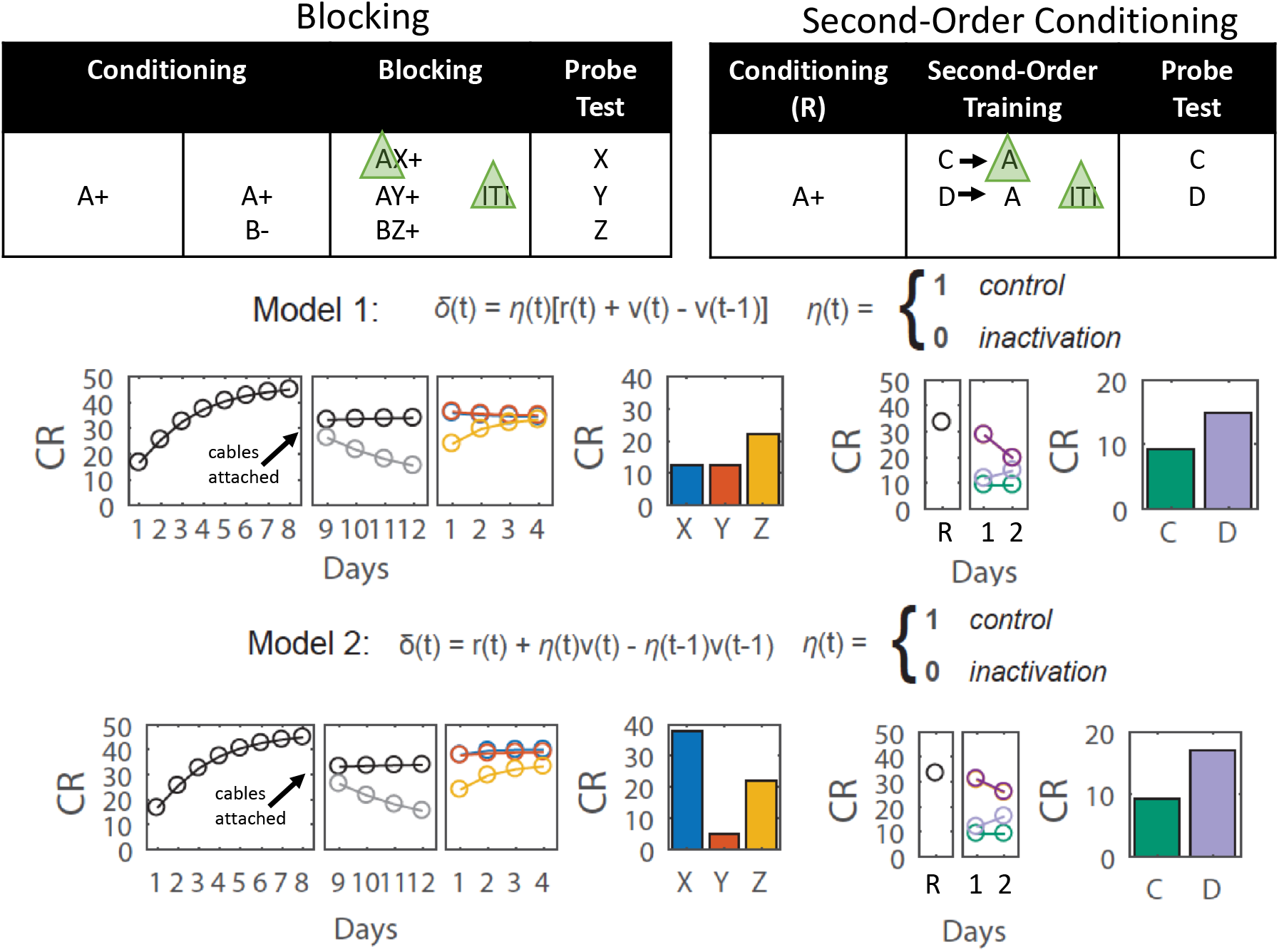
Experimental design for within-subjects blocking and second-order conditioning as used in our study, along with graphs modeling the predicted results of shunting of the dopamine transient at the start of the reward-predictive cue, A, in each procedure. In Model 1 the VTA DA signal encodes a prediction error and in Model 2 it encodes a reward prediction (Model 2). Bar graphs are reproduced from Figure 1 in the main text; other panels model results of training in the other phases. Note the output of the classic TDRL model was converted from V to conditioned responding (CR) to better reflect the behavioral output actually measured in our experiments. The major impact of this was on responding to Y in Model 2. Elimination of the prediction on AX trials in this model causes a positive prediction error on reward delivery in the blocking phase. This results in unblocking of X, however it also causes additional conditioning of A. Because we also used A as the blocking cue for Y, the larger but unmet prediction of A on these trials causes Y to become a conditioned inhibitor. This is not something that can be shown effectively in behavior unless Y is paired with a novel conditioned excitor.

**Figure S2:**
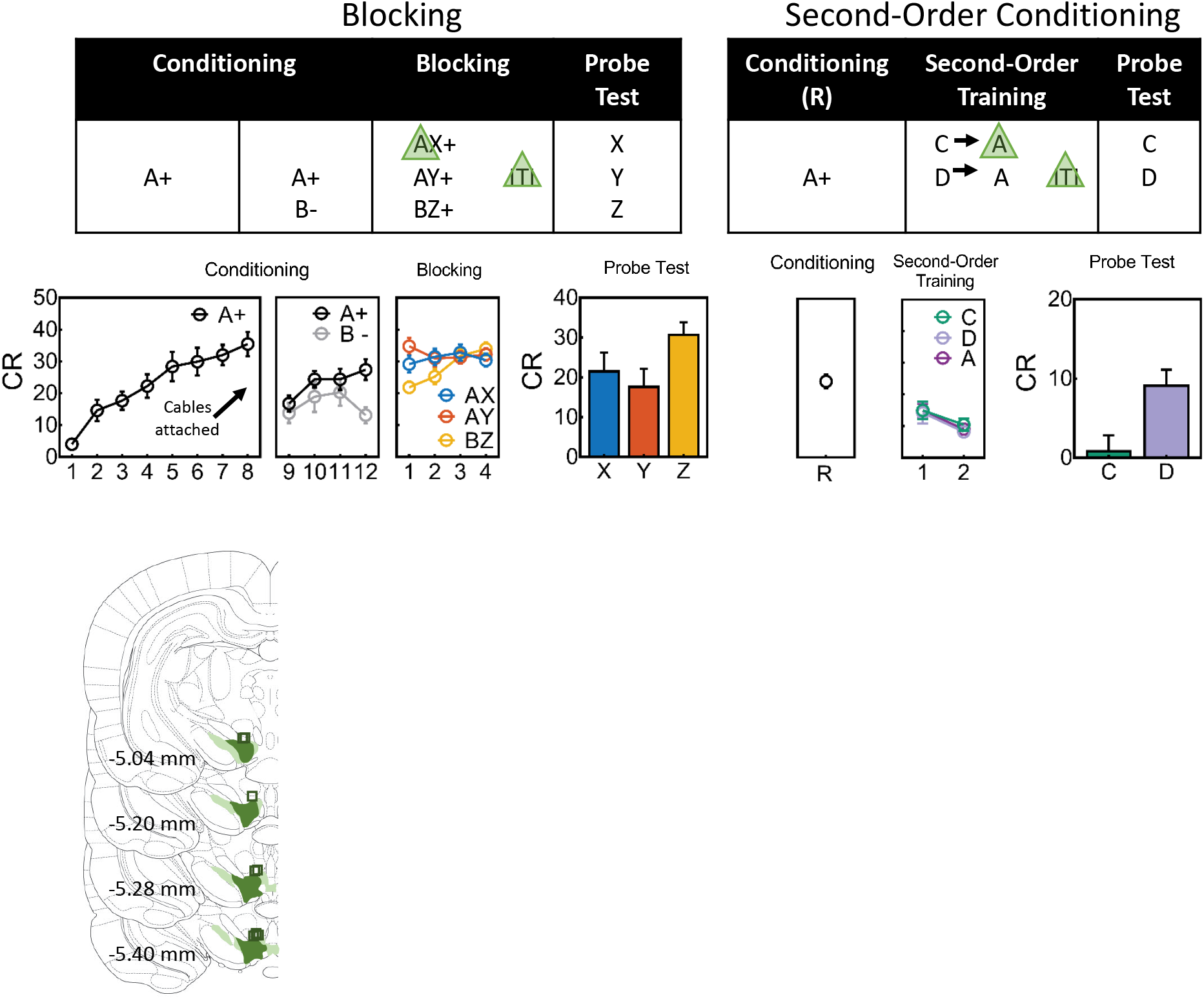
Experimental designs for within-subjects blocking and second-order conditioning as used in our study. During Conditioning responding to A (FI,15=21.8, Cl {0.42:1.13}) but not D (F<1, Cl {-0.41:0.39}) increased across days, and this responding was higher for A compared to D (FI,15=8.2, Cl {0.12:0.83}). During Blocking responding to the control compound (DZ) was lower compared to blocking compound (AX, AY) at the start of training (Dl: FI,15=12.9, Cl {-0.26:-1.00}) but equivalent by the end (D4: F<1, Cl {-0.67:0.30}), there were no differences between the blocking compounds (maxFl, 15=1.3, Cl {-1.01:.32}). Responding during the Probe test showed evidence of blocking (X and Y vs. Z) and no differences between the blocking cues (X vs Y, see Fig 3 legend for statistics). Responding to the retrained cue A increased across reminder (R) trials (FI,15=38.0, Cl {0.81:1.70}) while that to C (i.e., C→A trials) and D (i.e., D->A trials) did not differ across second-order conditioning (D1: F<1, Cl {-0.53:0.60}; D2: F<1, Cl {-0.30:0.56}). On Probe Test, responding to C was lower compared to D (see Fig 2 legend for statistics). Some data are reproduced from Figures 2 and 3 in the main text. CR or conditioned responding is percent time spent in the magazine during the last 5s of the cue. Drawings to the left illustrate the extent of expression of NpHR and location of fiber tips within VTA.

## Online Methods

### Modeling

Simulations of the behavioral designs were run using a one-step temporal difference learning algorithm, TD(0)^13^. This algorithm was used to estimate the value of different states of the behavioral paradigm with states being determined by the stimuli present at any particular time. Linear function approximation was used in order to estimate the value, V, of a given state, *s_t_*, by the features present during that state according to

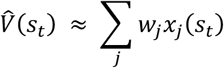

where *j* is indexed through all possible components of the feature vector *x* and corresponding weight vector *w*. The feature vector is considered to be the set of possible observed stimuli such that if stimulus *j* is present during state *s* at time *t*, then *x_j_*(*s_t_*) = 1, and zero otherwise. The weights are adjusted over time in order to best approximate the value of each state given the current set of stimuli. Weights, *w_j_*, corresponding to each feature, *x_j_*, are updated at each time step according to the TD error rule

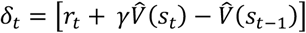

under linear value function approximation where γ is the temporal discouting factor. The weights are updated as

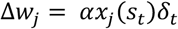

in which the scalar α is the learning rate. The linear value approximation reduces the size of the possible state space by generalizing states based on the features present. This approximation results in the calculation of the total expected value of a state as the sum of the expected value of each stimulus element present in the current state, a computation which is consistent with a global prediction error as stipulated by the Rescorla-Wagner model ^13^.

### Modeling of optogenetic manipulation of midbrain dopamine activity

Optogenetic inhibition of dopaminergic neurons was modeled two different ways to align with the different hypotheses of dopamine function described in the main text.

#### Model 1: Dopamine transients correspond to TD errors

For this model, inhibition of dopaminergic activity disrupts solely the error signal^10^

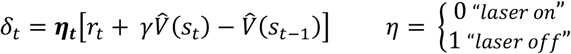

where *η* is a binary value determining whether the inhibition was present or not during state *s_t_*.

#### Model 2: Dopamine transients correspond to expected value

In this case, the dopaminergic inhibition disrupts the future expected value during the current state, and, since this becomes the prior expected value in the next state, the inhibition disrupt this as well

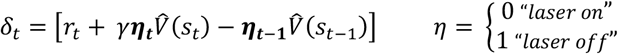

where *η* again determines whether the inhibition was present.

### Model Parameterization

Generalization of value across stimuli was modeled by setting the initial weights, *w_j_*, of a stimulus to 0.7 for stimuli of the same modality and 0.2 for stimuli of different modalities.

Conditioned responding to the food cup, *CR*, at each state was was modeled using a logistic function

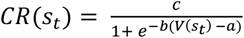

in which the parameters were determined based on empirical estimates of the maximal responding, *c*, the baseline responding, *a*, as well as the steepness of the learning curve, *b*. These were set as 55, 0.4, and 3 respectively for all simulations. Reduced responding to the foodcup while rats were attached to the patch cables was modeled as a reduction in the maximal responding to 40.

All simulations were performed with α = 0.05 and γ = 0.95. To ensure that order of cue presentations did not affect the findings, cue presentations during each stage of conditioning were pseudo-randomized and results of the simulations were averaged over 100 repetitions of the model. Simulations were performed using custom-written functions in MATLAB (Mathworks, Natick, MA), which are posted on Github (https://github.com/mphgardner/Basic_Pavlovian_TDRL/tree/Maes_2018).

### Subjects

Sixteen experimentally naïve Long-Evans transgenic rats (male: *n*=9, 390-587; female: *n*=7, 302-370g) expressing Cre recombinase under control of the tyrosine hydroxylase promoter (Th-cre+/-) were used for this experiment. The rats were bred inhouse (NIDA animals breeding facility) and implanted with bilateral optical fibers in the ventral tegmental area (VTA) at approximately 4 months of age.

### Surgical Procedures

Surgical procedures have been described elsewhere^8,9^. Rats were infused bilaterally with 1.2μL AAV5-EF1α-DIO, eNpHR3.0-eYFP into the VTA at the following coordinates relative to bregma: AP: −5.3mm; ML: ±0.7mm; DV: −6.55mm and −7.7 (females) or −7.0mm and −8.2mm (males). The viral vector was obtained from the Vector Core at University of North Carolina at Chapel Hill (UNC Vector Core). During surgery, ferrules carrying optical fibers were implanted bilaterally (200μm diameter, Precision Fiber Products, CA) at the following coordinates relative to bregma: AP: −5.3mm; ML: ±2.61mm, and DV: −7.05mm (female) or −7.55mm (male) at an angle of 15° pointed toward the midline.

### Apparatus

Experiments were conducted in 8 behavioral chambers (Coulbourn Instruments; Allentown, PA), which were individually housed in light- and sound-attenuating boxes (Jim Garmon, JHU Psychology Machine Shop). Each chamber was equipped with a pellet dispenser that delivered 45-mg sucrose pellets into a recessed magazine when activated. Access to the magazine was detected by means of infrared detectors mounted across the opening of the recess. Two differently shaped panel lights were located on the right-hand wall of the chamber above the magazine. The chambers contained a speaker connected to white noise and tone generators and a relay that delivered a 5kHz clicker stimulus. A computer equipped with GS3 software (Coulbourn Instruments, Allentown, PA) controlled the equipment and recorded the responses. Raw data were output and processed in Matlab (Mathworks, Natick, MA) to extract relevant response measures, which were analyzed in SPSS software (IBM analytics, Concordia University license, Canada).

### Housing

Rats were housed singly and maintained on a 12-hour light-dark cycle, where all behavioral experiments took place during the light cycle. Rats had *ad libitum* access to food and water unless undergoing behavioral testing, during which they received sufficient chow to maintain them at ~85% of their free-feeding body weight. All experimental procedures were conducted at the NIDA-IRP, in accordance with Institutional Animal Care and Use Committee of the US National Institute of Health guidelines.

### General behavioral procedures

Trials consisted of 13s visual and 10s auditory cues as described below; visual cues were 3s longer to allow optogenetic manipulation of the dopamine transient at the start of visual cue A without any interference with processing of other cues. Trial types were interleaved in miniblocks, with the specific order unique to each rat and counterbalanced across groups. Intertrial intervals varied around a 6-min mean (4-8 min range). All rats were trained during the light cycle between 10am and 4pm. Five 10s auditory (tone, clicker, white noise for X, Y, Z in blocking; chime and siren for C and D in second-order conditioning) and two 13s visual stimuli (flashing light and steady light for A and B) were used. The stimuli were counterbalanced across rats within each modality, and the reward used throughout consisted of two 45mg grape-flavoured sucrose pellets (5TUT; TestDiet, MO).

Training consisted of three phases: Conditioning, Compound Conditioning, and Test.

### Conditioning

Conditioning took place across 12 days (8 untethered days, 4 tethered days) and each day consisted of 14 presentations of A→2US, where a 13s presentation of A was immediately followed by two 45mg sucrose pellets (5TUT; TestDiet, MO). Towards the end of Conditioning (on tethered days 9-12), the rats also received four trials per day of non-reinforced presentations of B. This was done to reduce unconditioned orienting to the novel visual stimulus that would detract from learning on the first few trials of the compound stimulus ^15^. Responding (Figure 3 and Figure S2: Conditioning) to A (F_1,15_=21.8, CI {0.42:1.13}) but not B (F<1, CI {-0.41:0.39}) increased across days, and this responding was higher for A compared to B (F_1,15_=8.2, CI {0.12:0.83})

### Blocking

Following Conditioning, all rats received four days of Compound Conditioning, that is Blocking. Blocking followed the initial Conditoning phase because it was paramount that the reinforced cue had not been experienced in the absence of reiforcement (as in second order conditioning) as this could compromise its effectivenss to blocking learning. During this phase two compounds consisting of the pre-trained cue A and a novel auditory cue, X or Y, and a third compound consisting of the pre-exposed cue B and a novel auditory cue Z were presented. Each compound received six reinforced trials with the same reward (AX→2US; AY →2US; BZ→2US). This yielded two blocking compounds AX, AY, and a control compound, BZ. The presentation of the 13s visual cues began 3s prior to onset of the 10s auditory cues. On AX trials, continuous laser light (532nm, 18-20mW output, Shanghai Laser & Optics Century Co., Ltd) was delivered into the VTA for 2.5s. starting 0.5s prior to the onset of A; on AY trials, the same light pattern was delivered during the intertrial interval, 120-180s after termination of the compound. During blocking (Figure 3 and S2: Blocking), responding to the control compound (BZ) was lower compared to that seen to the blocking compounds (AX and AY) on the first day of training (F_1,15_=12.9, CI {-0.26:-1.00}) but similar on subsequent days (D2: F_1,15_ = 2.68, CI {-0.12:0.94}; D3: F<1, {-0.46:0.50}; D4: F<1, {-0.67:0.30}). Responding to the blocking compounds (AX and AY) did not differ across this phase of training (D1: F_1,15_ = 1.3, CI {1.00:0.32}; D2: F<1, {-0.64:0.70}; D3: F<1, {-0.43:0.62}; D4: {-0.64:0.38}). The lack of differences between AX and AY provide evidence that shunting DA firing during the start of A did not disrupt processing of A.

To confirm learning and determine the effect of stimulation of TH+ neurons in the VTA, rats received a probe test in which each of the auditory cues (X, Y, and Z) was presented four times alone and without reward for a total of 12 trials. Rats received the same probe test again two days afterwards, which allowed for behavioural recovery. The two test sessions were collapsed. Analyses focused on the pooled data from the initial trial in each test. The first trial eliminates any within-session effects of non-rienforcement, which can mask behavioural differences.

### Second Order Conditioning

Prior to start of Second Order Conditioning, all rats received a single reminder session for A, which consisted of retraining of the A→2US contingency across 14 trials as described above. This was done to offset any effects of probe testing without reward at the end of blocking. Magazine responding to the retrained cue A (Figure 2 and Figure S2) increased across trials (F_1,15_=38.0, CI {0.81:1.70}). Following retraining, rats received two sessions of second-order conditioning consisting of six presentations of X and six presentations of Y each paired with A (C→A; D→A). On C→A trials, continuous laser light 532nm, 18-20mW output, Shanghai Laser & Optics Century Co., Ltd) was delivered into the VTA at the start of A in the same manner as that used in Blocking; on D→A trials, the same light pattern was delivered during the intertrial interval, 120-180s after termination of A. Responding to C and D (see Figure 2 and S2: Second-order training) did not differ across second-order conditioning (D1: F<1, CI {-0.53:0.60}; D2: F<1, CI {-0.30:0.56}), there were no effects of trials (D1: F<1, CI {-0.27:0.36}; D2: F<1, CI {-0.59:0.63}) and no interaction (D1: F<, CI {-0.53:0.53}; D2: F_1,15_=1.7, CI {-0.18:0.77}). Similarly, responding to A following C or D did not differ (D1: F_1,15_=1.9, CI {-0.14:0.67}; D2: F_1,15_=1.1, CI {-0.27:0.81}), there was no effect of trials (D1: F<1, CI {-0.41:0.36}; D2: F<1, CI {-0.52:0.58}), nor an interaction (D1: F_1,15_=1.5, CI {-0.46:0.13}; D2: F<1, CI {-0.30:0.81}). Therefore, responding to A was combined. Following this training, rats received a probe test where cue C and D were each presented six times in the absence of any reinforcement (Figure 2 and S2: Probe Test).

As mentioned in the main text, the differential effects of VTA DA shunting during A in second-order conditioning and blocking provide evidence that the disruptive effect on the former is not due to light artefacts serving to hinder processing of A. If so, then, we would see a disupriton of the blocking effect as well (i.e., learning about X). These results support temporal difference accounts by providing causal evidence that cue-evoked dopamine transients function as prediction errors.

### Histology

All rats were euthanized with an overdose of carbon dioxide and perfused with phosphate buffered saline (PBS) followed by 4% Paraformaldehyde (Santa Cruz Biotechnology Inc., CA). Fixed brains were cut in 40μm sections, images of these brain slices were acquired and examined under a fluorescence (Olympus Microscopy, Japan). The viral spread and optical fiber placement was verified and later analyzed and graphed using Adobe Photoshop.

